# Novel tuberculosis combination regimens of two and three-months therapy duration

**DOI:** 10.1101/2022.03.13.484155

**Authors:** Tawanda Gumbo, Moti Chapagain, Gesham Magombedze, Shashikant Srivastava, Devyani Deshpande, Jotam G. Pasipanodya, Ruramai Gumbo, Keertan Dheda, Andreas Diacon, David Hermann, Debra Hanna

## Abstract

The tuberculosis treatment duration is 4-6 months, but can be 18–20 months with multidrug resistance. Ultrashort regimens that cure regardless of resistance status [“pan-tuberculosis regimens”] would be a major step towards global tuberculosis control. Starting with seven drugs [bedaquiline, delamanid, pretomanid, OPC-167832, sutezolid, moxifloxacin, and pyrazinamide] in clinical testing, we calculated that there were thousands of possible novel combinations to evaluate. We used mathematical and pharmacokinetics/pharmacodynamic-based hollow fiber modeling to reduce this complexity to nine combinations, which we tested. Each regimen’s fast and slow growth bacteria kill-slope trajectories and times-to-extinction were estimated and translated to minimum required therapy duration in patients. Sputum microbial kill trajectories of two combination regimens that have undergone clinical testing were correctly predicted by this approach. Four pan-tuberculosis regimens were predicted to achieve relapse-free cure after 2-3 months therapy in patients. The kill-slopes were used to design expedited clinical trials that minimize patients’ risks.

## Introduction

Tuberculosis [TB] causes 3% of all human deaths each year. Approximately 20% of the deaths are due to drug-resistant TB [DR-TB]: multi-drug resistant [MDR] TB [resistance to isoniazid and rifampicin] has a case-fatality rate of 40%.^1^ Fortunately, the discovery of new anti-TB agents such as bedaquiline, delamanid, pretomanid, OPC-167832, and sutezolid, has increased the possibility of designing combination regimens that could cure such DR-TB as easily as drug-susceptible TB, termed “pan-TB” regimens.^2-10^ Here, we chose bedaquiline [D], delamanid [D], pretomanid [Pa], OPC-167832 [O], sutezolid [S], moxifloxacin [M], and pyrazinamide [Z], for the *de novo* design of such regimens, since these drugs have undergone at least proof-of-concept clinical testing, have completed formal pharmacokinetics-pharmacodynamics [PK/PD] studies, have better kill rates compared to congeners, and have oral formulations.^2-15^ We hypothesized that “pan-TB” regimens based on three to four of these seven drugs could shorten therapy duration from the current six [short course] months to less than 4 months.

Historically, TB regimens were designed based on the order in which the drugs were discovered, leading to the current gold standard of rifampin [or rifapentine], isoniazid, pyrazinamide, ethambutol, and moxifloxacin for 4-6 months. However, at the time studies reported here were done, there were already several TB drugs to consider that could be combined to form hundreds of regimens. Thus, tools to identify specific promising regimens for rapid clinical trial evaluation were needed. One such tool we considered was the hollow fiber system model of TB [HFS-TB], with a quantitative forecasting accuracy of 94% for clinical response rates and optimal exposures.^16^ This tool has been formally qualified as a drug development tool by the European Medicines Agency^17^ and utilized to perform PK/PD monotherapy studies for microbial kill and resistance suppression for >50 individual drugs as well as to design several new combination regimens.^2-5,7,10,11,18,19^

In order to accurately translate therapy duration in the clinic from HFS-TB results, we developed ordinary differential equations [ODEs] that describe *Mycobacterium tuberculosis* [*Mtb*] kill trajectories of log-phase growth [“fast”] and semidormant/non-replicating persistent [NRP] “slow” bacilli on treatment with standard 6-months therapy; this relied on repetitive sampling of HFS-TB units and sputa of patients.^20,21^ We designated the fast replicating *Mtb* kill slope *γ*_*f*_ and that for the slow bacilli *γ*_*s*_. The ODEs were used to estimate the time-to-extinction [TTE] of the entire *Mtb* lung population [i.e, to *Mtb* burden 10^−2^ CFU/mL]^20^ based on the HFS-TB data, which was characterized by a highly homogeneous distribution of initial *Mtb* burden, and highly controlled *Mtb* growth conditions and drug pharmacokinetics, as one would expect from an engineered *in vitro* model.^20,21^ We also estimated patients’ sputa *γ*_*f*_, *γ*_*s*_, and TTE using the same ODEs, in clinical data characterized by a wide range of initial *Mtb* sputum burden, lesion heterogeneity, and pharmacokinetic variability, as is typical of patients.^20-22^ The two datasets formed a structure-preserving map for *γ*_*f*_, *γ*_*s*_, and TTE, that is a morphism, as defined by category theory.^23-25^ This meant that the two datasets could translate one to another using a non-linear multistep function. In category theory, the complexity of human TB lesions is not an impediment to identifying relationship functions with *in vitro* datasets, rather what is important is that the datasets map one to another based on identity functions. In fact, the TB lesion complexity and heterogeneity are why the translation factor we identified is complex, multi-step, and non-linear.^20^ For validation, we identified *Mtb γ*-slopes and TTE in 108 HFS-TB units treated with the REMoxTB experimental regimens, and used the translation platform to predict cure rates in 1,932 patients on the 4-months regimens, which were compared to observed outcomes in the actual 1,932 patients recruited to the REMoxTB clinical trial. The model predicted vs observed relapse-free cure rates were 77.0% [95% confidence interval (CI): 74.4%-79.6%] vs 77.7% [95% CI, 74.3%-80.9%] and 76.4% [95% CI, 73.9%-79.0%] vs 79.5% [95% CI, 76.1%-82.5%]) in the experimental arms.^20^ Our translation platform also predicted a 72% relapse-free cure on the 4-months duration PaMZ regimen versus the 71% recently documented in interim trial results.^26^ Here, we identified the *γ-*slopes and TTEs of novel combination regimens in the HFS-TB, translated them to patients, and ranked regimens based on the shortest therapy duration associated with relapse-free cure.

## RESULTS

### HFS-TB and modeling to reduce complexity to tractable possibilities

For just the seven TB drugs we studied, how many possible 2, 3, and 4-drug combination regimens were there? Two-drug combinations were for optimal design purposes only and were not meant for clinical use. **Figure 1** shows that based on Fermat and Pascal,^27^ there are 91 possible combinations, assuming that only one dose is used per drug. Based on Shannon entropy, optimal Fisher information [the most information-rich drug exposures (EC)] are expected to occur at exposures mediating 20% [EC_20_], 50% [EC_50_], and 80% [EC_80_] of maximal kill, and are the minimum that can be used for optimal design of full exposure-effect surface experiments to identify concentration-dependent synergy, or additivity, or antagonism.^28^ This calculates to 336 exposure-response units and 11,536 units/possibilities [**Figure 1]**. If two HFS-TB replicates were used for each then 23,072 units are required, or if 34 mice per regimen were used for relapse studies as in Tasneen et al’s experiments^6^ then 392, 224 mice would be required. These numbers of mice or HFS-TB units are intractable.

**Figure 1.**
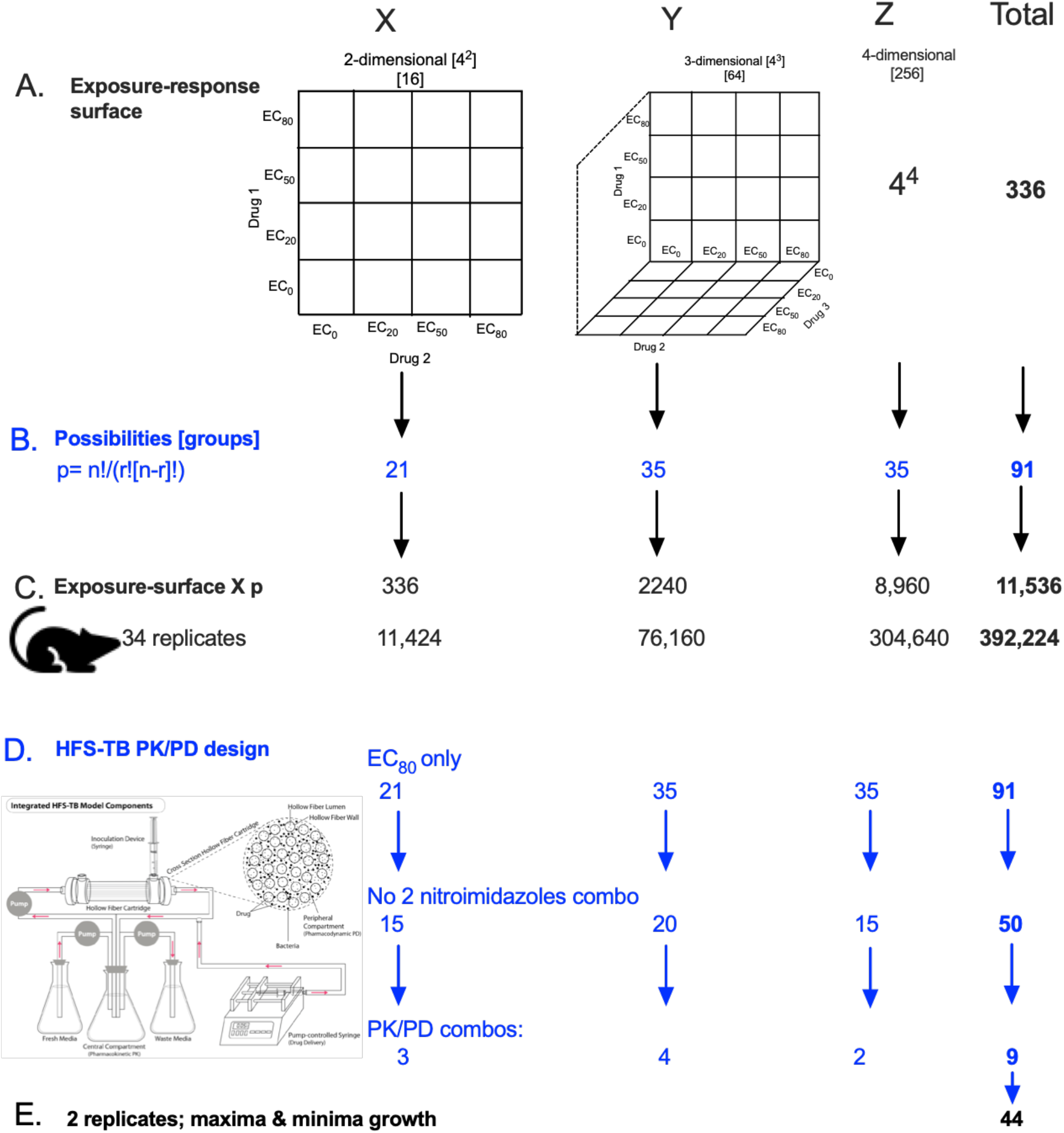
Optimizing design to reduce complexity. Increasing drug concentrations are depicted as vectors in Euclidian space, a 2-drug combination is a 2-dimensional surface, shown with data points for the four exposures that creates 4^r^ experimental groups, and generalized for r drugs in a regimen [r-dimensional surface] to 4^*r*^. Here we show r=2, r=3, and r=4, so that column X is for r=2, column Y for 3-drug regimen, and column Z for 4-drug regimens. The number of units in each row is shown in rows A-D. In row B, the total number of possibilities [P] of combinations with 7 drugs [n=7], for *r-*drug combination regimen, are shown. However, since each combination requires examination of the whole exposure surface [defined by at least 4 exposures], the number of possibilities increases as shown in row C. We consider the example of the mouse TB relapse model assuming a total of 34 mice per regimen (*3*). In Row D, if only the HFS-TB derived EC_80_ is used, and we disallowed combinations of the two nitroimidazoles, and discarded combinations that have demonstrated meager improvement over standard therapy in prior HFS-TB PK/PD work [e.g., moxifloxacin plus either pyrazinamide or oxazolidinone, oxazolidinone plus pyrazinamide, oxazolidinone plus nitroimidazole, while moxifloxacin-pyrazinamide-oxazolidinone combination, among several, are not better than standard therapy], decreases the number of HFS-TB units, making the approach more tractable.

Since multiple [but not all] HFS-TB and clinical studies have shown that antagonism is encountered mostly at low concentrations and at EC_20_ and EC_50_ exposures while EC_80_s have generally been additive or synergistic,^22,29-32^ we reduced the possible combinations by examining only the EC_80_ in combination therapy in **Figure 1D**. Further PK/PD considerations shown in **Figure 1D**, and our prior work with multiple combinations, reduced the number of possibilities to the nine novel regimens in supplementary **Table S1**. Moreover, since the HFS-TB allows repetitive sampling at multiple points within the same system, and the main outcome was *γ*-slopes, two HFS-TB replicates could be used, leading to 22 HFS-TB units, including positive and negative controls. Extra complexity is added by the consideration that *Mtb* exists in human TB lesions under different stressors and growth rates, introducing a large spectrum of anti-TB therapy refractoriness from highly susceptible fast-to refractory slow-Mtb.^33-35^ We chose the two replication and therapy refractoriness extrema to test the 11 regimens in the HFS-TB for slow replicating [HFS-TB-slow] and fast replicating [HFS-TB-fast] *Mtb*.

### HFS-TB concentrations achieved in patient lesions and modeled to clinical doses

We compared the kill rates of the eleven regimens in the HFS-TB-fast and HFS-TB-slow, as described in full in the online supplement. Since for each drug the area under the concentration-time curve [AUC_0-24_] to minimum inhibitory concentration [MIC] is the PK/PD driver, except pretomanid [which time-driven, but nevertheless has a half-life of 16-19hrs],^2-15^ we administered each drug on a once-a-day dosing schedule in the HFS-TB combination regimens. We measured drug concentrations at nine time-points over 72 hours, and the concentration-time profiles demonstrated exposures shown in **Figure 2**. Since the most highly ranked determinants of microbial outcomes in patients with TB are drug concentrations, between-patient pharmacokinetic variability, *Mtb* MIC variability, PK/PD exposures, and lung intralesional penetration, we integrated each of these factors in Monte Carlo experiments to identify clinical doses that would achieve or exceed the HFS-TB exposures shown in **Figure 2**, using steps outlined in online methods.^2,4,7,10,22,32,36-52^ **Figure 3** and supplementary data text show that the *in-silico* dosing achieved the same serum pharmacokinetic parameters and concentrations observed in actual patients. **Figure 4** shows that the bedaquiline, delamanid, OPC, pretomanid, and sutezolid exposures achieved in our HFS-TB experiments could easily be achieved in lung TB cavities at doses tolerated by patients in the clinic. However, the pyrazinamide and moxifloxacin doses identified were very high and might not be tolerated by patients, while rifampin and isoniazid doses were also high but in the range that could be tolerated by patients.^39,53^ Thus, the first output of our approach was not just which drugs to combine in regimens, but also the clinical doses to combine.

**Figure 2.**
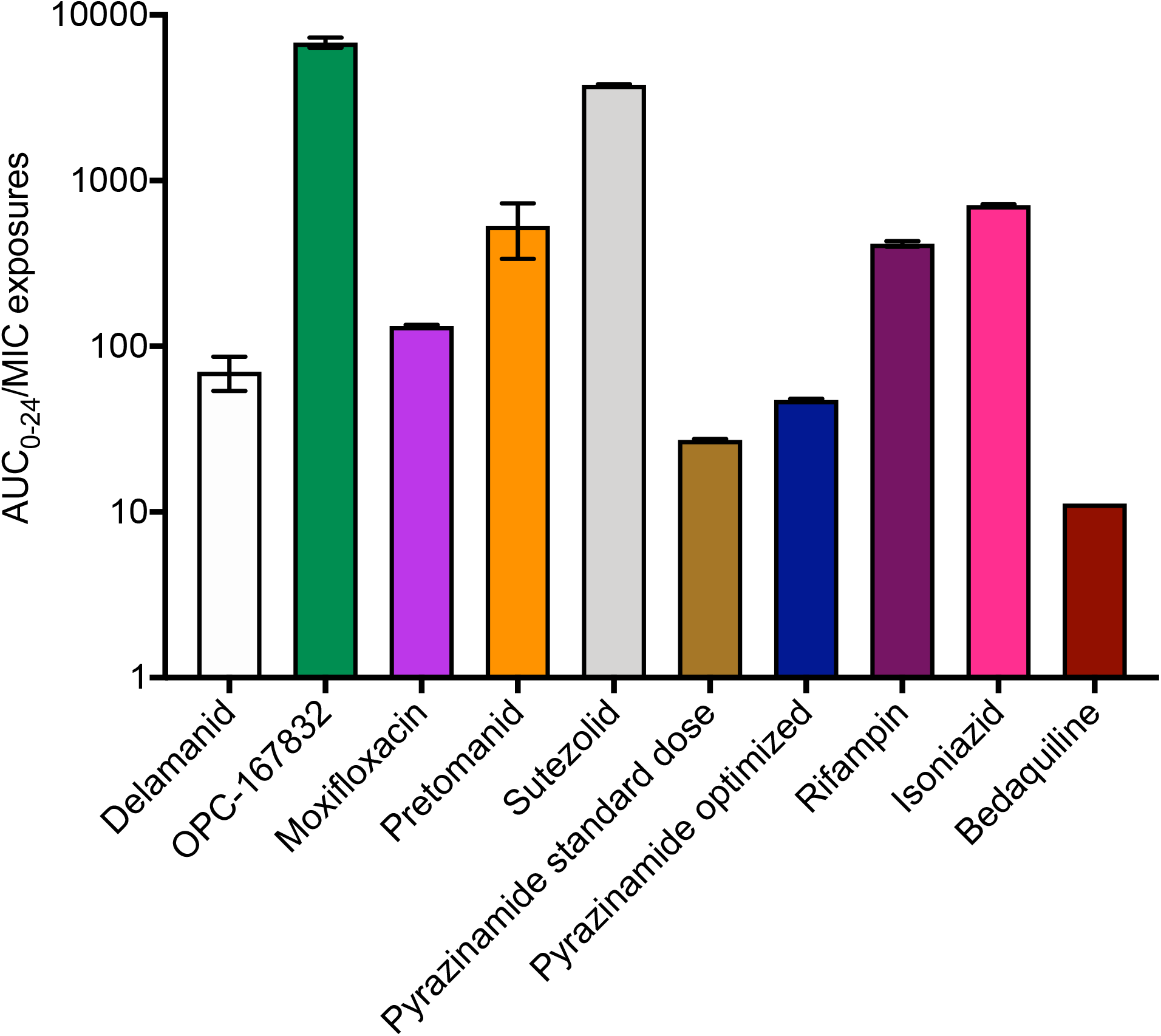
Exposures achieved in the HFS-TB. The bar graph shows mean values of all HFS-TB replicates while the error bars are standard deviation. Since all the drugs, except pretomanid, have PK/PD driver of AUC_0-24_/MIC, the drug exposures calculated from the measured drug concentrations in the HFS-TB are expressed as AUC/MIC ratios. The pretomanid % time that concentration persists above the MIC was 100%. The exposures identified in the HFS-TB are equivalent to those achieved at site of infection; the error bars show minimal technical variability between HFS-TB units. For OPC, the exposures were adjusted for accumulation of the drug in the system. The results show that the intended EC_80_s were achieved for each drug and that for standard dose for bedaquiline.

**Figure 3.**
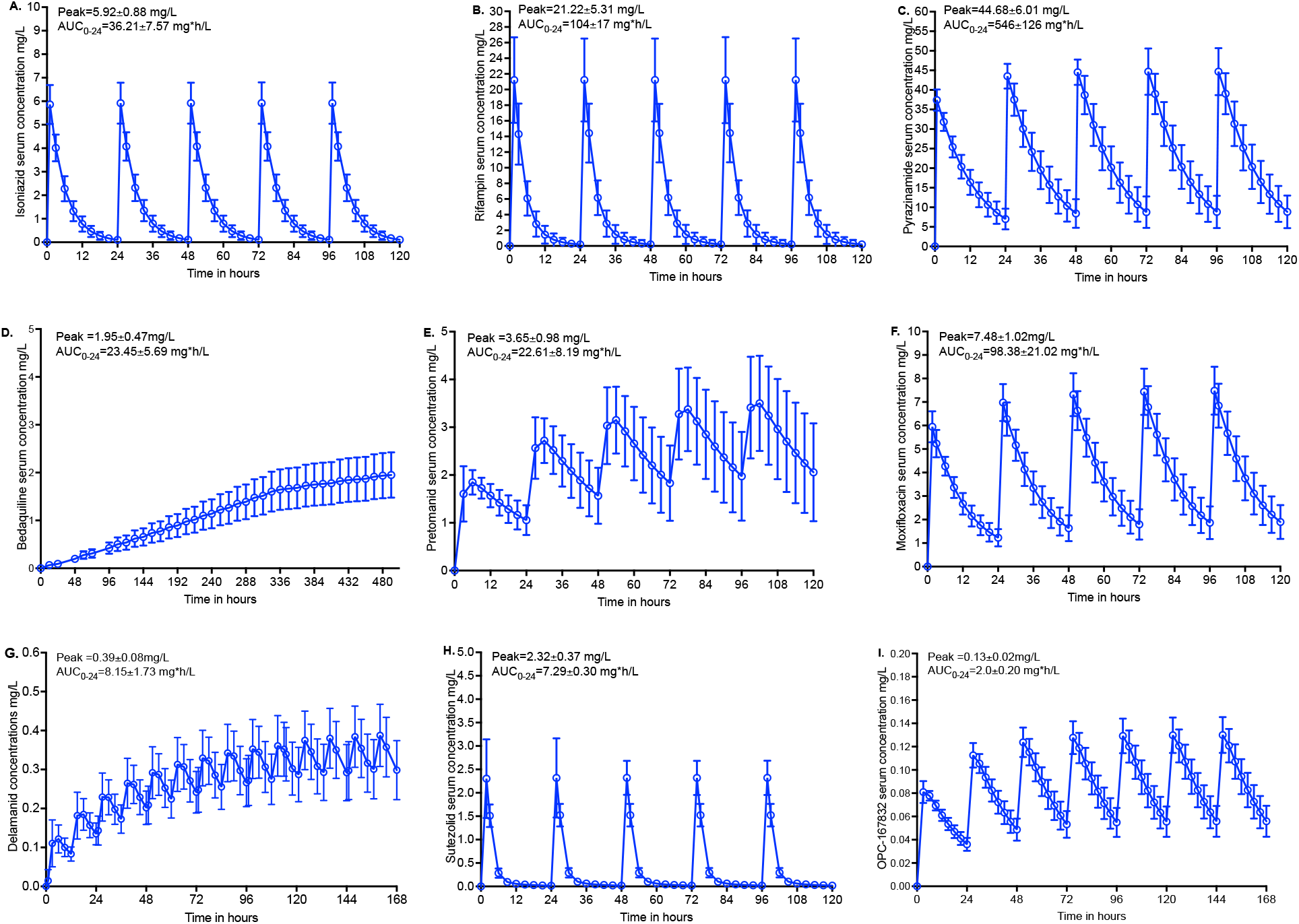
Pharmacokinetic parameters of nine drugs in Monte Carlo experiments of 10,000 patients. Symbols are mean concentrations and error is standard deviation. The AUC_0-24_s were then transformed into lung intralesional concentrations based on penetration ratios shown in the methods section. **A**. Isoniazid 600 mg/day. **B**. Rifampin 35 mg/kg/day. **C**. Pyrazinamide 2,000 mg/day. **D**. Bedaquiline dosing simulated was 400 mg a day for 2 weeks, followed by 200 mg three times a week, thus the “steady” state period stretches out to 3 weeks. **E**. Pretomanid. **F**. Moxifloxacin. **G**. Delamanid 100 mg twice a day. **H**. Sutezolid. **I**. OPC-167832.

**Figure 4.**
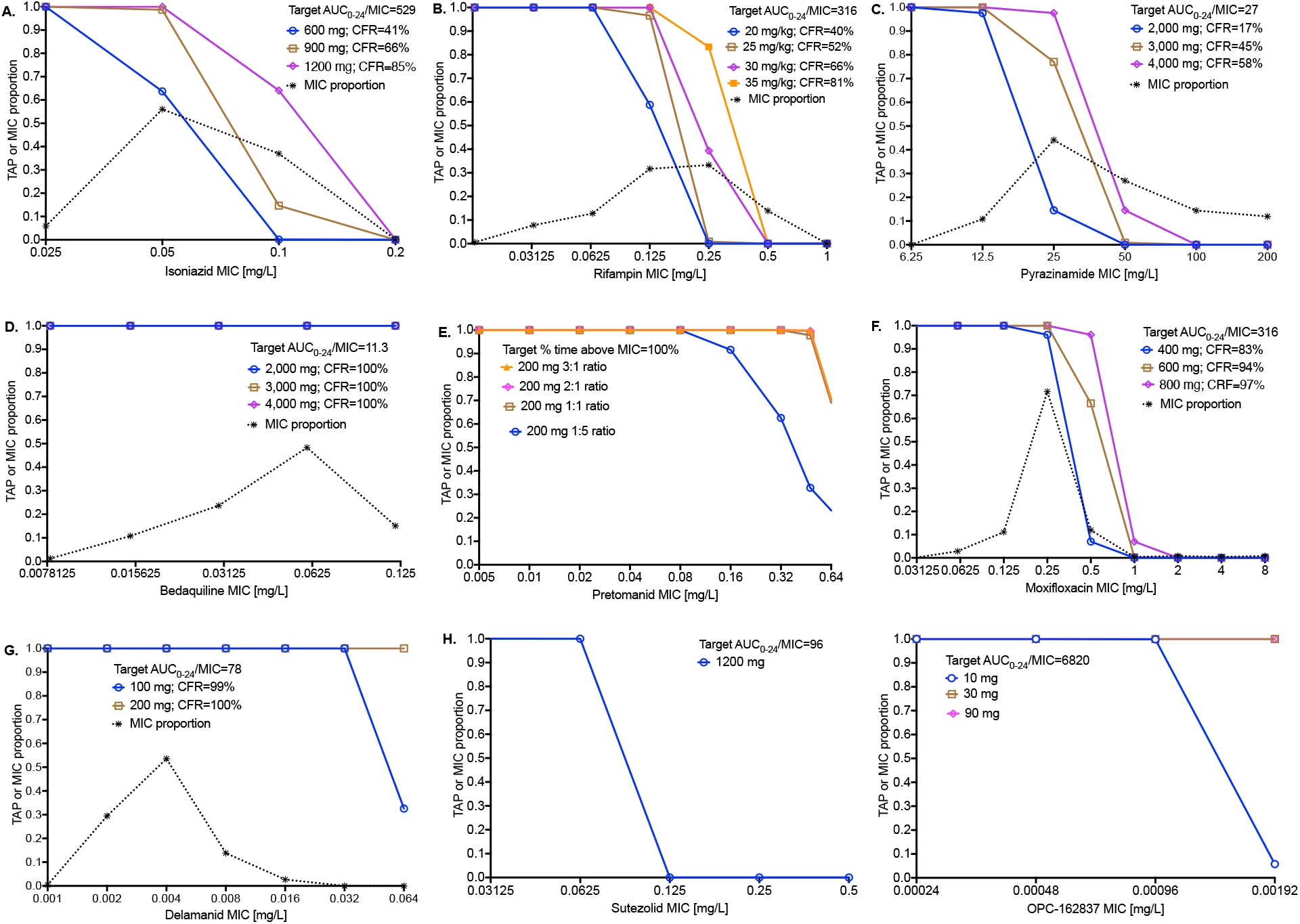
In silico dose-finding in 10,000 simulated subjects. All doses are per day. The intralesional target exposure is the exposure achieved in HFS-TB shown for each panel. Target attainment probability [TAP] is the proportion of patients treated with a dose, who achieve, or exceed, the target exposure in TB lung cavities, based on serum-to-caseum penetration ratios, at each MIC *I*. The cumulative fraction of response [CFR] is the proportion of all patients who achieve target attainment: 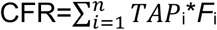, where *F*_*i*_ is the fraction of isolates at MIC_*i*_. **A**. Isoniazid >600 mg was required for good CFR. **B**. Rifampicin doses of 35 mg/kg were required for good CFR. **C**. The pyrazinamide CFR for 4G was only 58%. **D**. Bedaquiline easily achieved the target in TB lesions in all patients, even at 200 mg. **E**. For pretomanid, the standard dose of 200 mg would achieve % time above MIC of 100% if lung-to-serum penetration ratio was ≥1:1, but would fail if penetration was ≤20%. **F**. Moxifloxacin 800 mg achieved a CFR of 97%. **G**. Even though our PK simulation had underestimated the delamanid AUCs encountered in patients by 50%, the CFR was 99% on 100 mg. **H**. Sutezolid MIC distribution is poorly documented, with 6/7 isolates having MICs ≤0.0625 mg/L, making the CFR of 1200 mg of 86% imprecise. **I**. For OPC, the dose of 10 mg achieved maximal effect at all MICs in the range except the highest MIC. Thus 10 mg is likely as effect as higher doses.

### Ranking of regimens performance in the HFS-TB using traditional culture readouts Figure 5A

shows regimens’ performance in the HFS-TB-slow, based on CFU/mL readout on Middlebrook agar. First, the delamanid-bedaquiline [DB] dual regimen demonstrated no effect. However, with the addition of OPC-167832 the DBO regimen became one of the most highly active, suggesting that addition of a third drug makes this dual combination an effective regimen. Second, the standard regimen performed as has been seen in the past in the HFS-TB and took 28 days to negative cultures on agar, a necessary quantity control step. Third, the BPaMZ regimen killed the entire slow *Mtb* population fastest. Fourth, **Figure 5A** shows that the second highest performing cluster of regimens, which eradicated *Mtb* by day 14 was comprised of five regimens: DBOS, DBO, DOS, BO and optimized PaMZ. In **Figure 5B**, Mycobacteria Growth Indicator Tube [MGIT] liquid culture time-to-positivity [TTP] readout, which is more sensitive than agar, was particularly useful in temporal resolution of bacterial eradication at the limits of CFU quantitation in this cluster of regimens: the DBOS, BOS, and high-dose moxifloxacin-and pyrazinamide-optimized PaMZ, had truly negative cultures by day 14, while DBO and BO dual combinations only achieved negative cultures on day 21. The ranking order of all regimens is shown in **Table 1**.

**Table 1.** Ranking of regimens in the HFS-TB using different measures of efficacy.

| Sterilizing effect: time to negative culture |  | Bactericidal effect: time to negative culture |  | Extended EBA log<sub>10</sub> cfu/ml/day |  | Extended EBA TTP [day/day] |  |
| --- | --- | --- | --- | --- | --- | --- | --- |
| Rank | Regimen | Rank | Regimen | Rank | Regimen | Rank | Regimen |
| 1 | BPaMZ | 1 | DBO | 1 | DOS | 1 | DBOS |
| 2 | DBOS | 2 | DBOS | 1 | DBOS | 1 | DO |
| 2 | PaMZ | 2 | DO | 1 | **Standard** | 1 | DBO |
| 2 | DOS | 4 | DOS | 1 | BO | 1 | DOS |
| 5 | BO | 4 | **Standard** | 5 | BPaMZ | 5 | BO |
| 5 | DBO | 6 | BO | 5 | DO | 6 | **Standard** |
| 7 | DO | 6 | Optimized PaMZ | 5 | PaMZ | 7 | PaMZ |
| 8 | **Standard** | 8 | BPaMZ | 5 | DBO | 8 | BPaMZ |
| 9 | BPaZ | 9 | BPaZ | 5 | BPaZ | 9 | BPaZ |
| 10 | BD | 10 | DB | 10 | DB | 10 | DB |

Ranking [starting with fastest kill is shown for microbial kill of slow replicating bacteria [sterilizing activity]. Microbial kill of fast replicating bacteria [bactericidal effect] is shown using different definitions of outcome and readouts, leading to changes in ranking of combination regimens.

**Figure 5.**
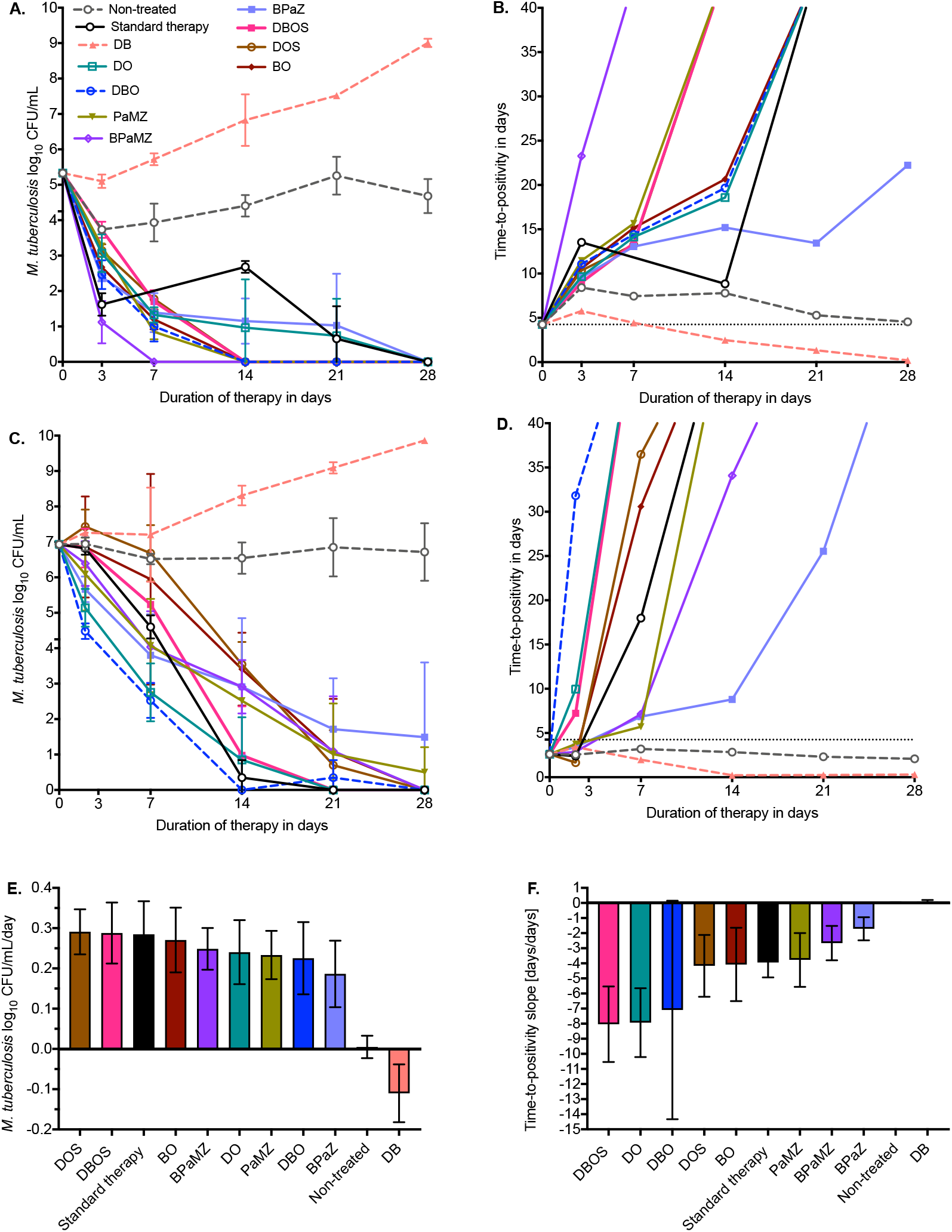
Bacillary subpopulation regimen kill trajectories in hollow fiber systems. Mean values and error bars [standard deviation] are shown based on colony forming units per mL [CFU/mL] and time to positivity [TTP] for two replicates. TTP is capped as negative after 43 days, making it difficult to calculate scatter and mean estimates when replicates have values >43 days. **A** and **B** are the sterilizing activity experiments, which show that the BPaMZ regimen was highest ranked based on shortest time to wipe out bacteria. **C-D** are bactericidal activity experiments: DBO was the highest ranked based on earliest time to negative cultures. **E** and **F** are 14-day bactericidal activity slopes, using log_10_ CFU/mL/day and TTP [day/day], leading to different rankings.

In the HFS-TB-fast, DB also demonstrated poor effect [**Figure 3C**]. However, the addition of OPC totally reversed that, and the DBO regimen became the highest ranked regimen of all by both CFU/mL [**Figure 5C**] and TTP [**Figure 5D**]. The second highest ranked group included DO and DBOS, which based on TTP killed faster than standard therapy. TTP demonstrated that standard therapy killed all *Mtb* by day 21, a performance similar to DOS and PaMZ. Since extended [14-day] early bactericidal activity [EBA_14_] is measured by comparing linear-regression-based kill slopes in the clinic, we identified EBA_14_ slopes from the HFS-TB-fast, as shown in **Figure 5E-F**. In patients on standard therapy that included a rifampin dose-finding study by Boeree et al,^39^ rifampin 35 mg/kg plus isoniazid and pyrazinamide with the same observed AUCs and AUC_0-24_/MIC as were attained in the HFS-TB [**Figure 2**], the EBA_14_ was 0.26 [95% CI: 0.19-0.34] log_10_ CFU/mL/day, similar to the 0.28±0.04 log_10_ CFU/mL in the HFS-TB-fast [**Figure 5E]**. Thus, our HFS-TB bactericidal activity results accurately reflected combination regimen EBA_14_ values in the clinic. However, **Figure 5C-F** and **Table 1** demonstrate that the ranking of regimens was different based on whether time to negative cultures [**Figure 5C-D**] or EBA_14_ was used [**Figure 5E-F**] for decision-making. In addition, there was also a difference in ranking between EBA_14_ by TTP versus log_10_ CFU/mL/day-based slopes.

### Analysis of HFS-TB results using ODEs, *γ*-slopes, and time to extinction

Since regimen effect in HFS-TB-fast and HFS-TB-slow ranked regimens differently but patients harbor both populations, and ranking depended on type of culture assay agar, and the kill slopes were clearly non-linear and therefore should not be analyzed using linear regression, we used our ODEs [incorporating both TTP and CFU/mL readouts] that simultaneously follow the trajectories of the fast-and slow-replicating *Mtb* bacterial subpopulations to identify *γ*_*f*_, *γ*_*s*_, and time-to-extinction [TTE] of all bacteria. The trajectories and TTEs defined by the ODEs are shown in **Figure 6** and tabulated in **supplementary Table S2. Supplementary Figure S1** shows the modeled growth of bacilli in non-treated controls and DB dual combination, which did not achieve bacterial population extinction. **Figure 6A** shows that with standard therapy, the driver of TTE was the *γ*_*s*_ slope, consistent with the current understanding of the special *Mtb* populations hypothesis and our models of sputa data.^33,34,54^ However, **Figure 6** also shows that for several experimental regimens either the 95% credible intervals of *γ*_*s*_ and *γ*_*f*_ overlapped, or in some cases the kill trajectories swapped positions and the *γ*_*f*_ took reach extinction than *γ*_*s*_. **Figure 6** shows that as regimens killed faster than standard therapy and TTEs became shorter and shorter, the *γ*_*f*_ *-*slope contribution to the TTE increased.

**Figure 6.**
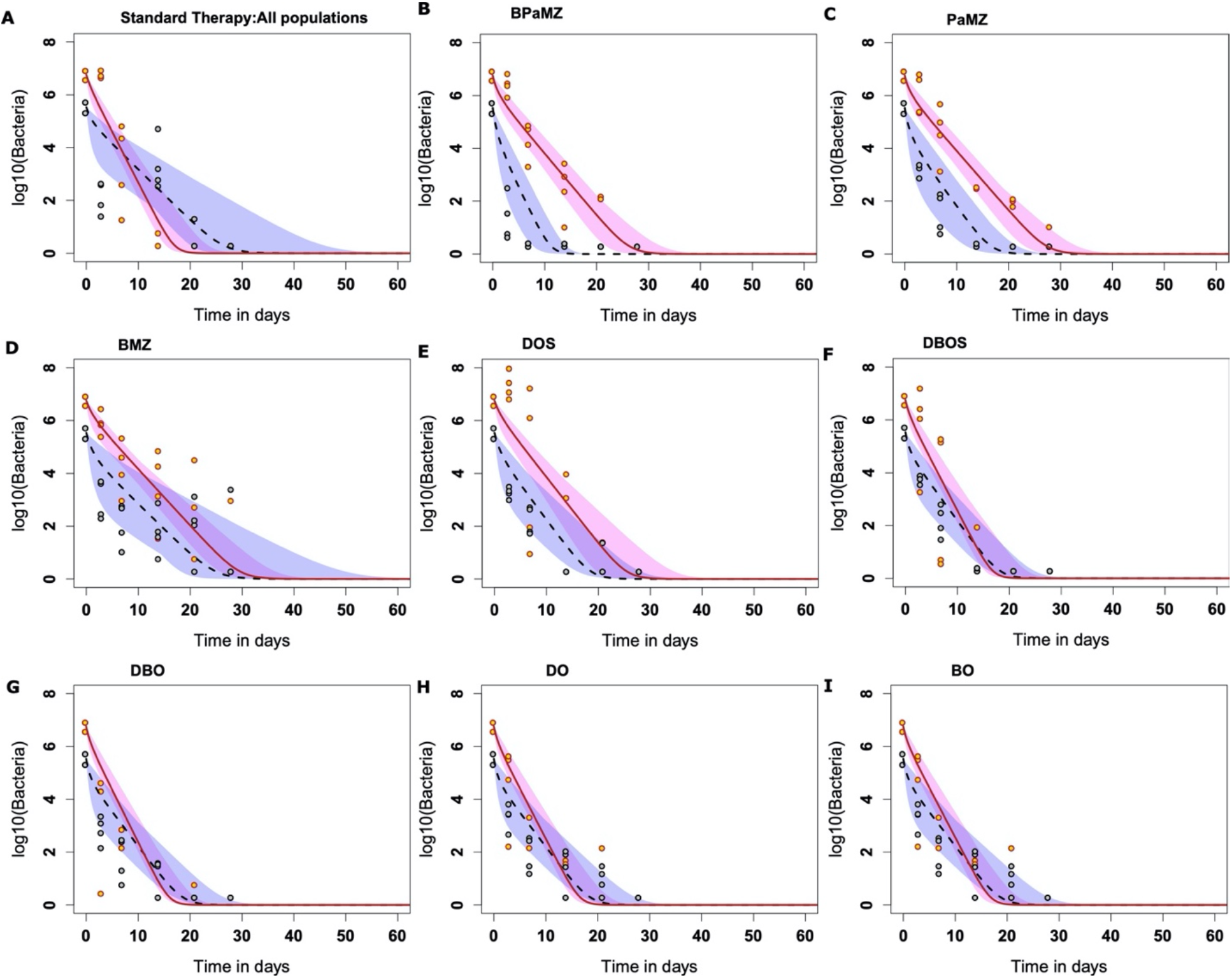
Bacillary subpopulation growth and regimen kill trajectories in the HFS-TB. *Mtb* burden changes with time [i.e., slopes] achieved by different drug combinations are shown in periwinkle for non/slow replicating [*γ*_*s*_] and in pink for fast replicating bacilli [*γ*_*f*_], until both populations reach extinction. The shaded regions represent 95% credible intervals for *γ*_*s*_ and *γ*_*f*_. (**A**) *Mtb* kill kinetics achieved by standard therapy demonstrate that the time-to-extinction [to *Mtb* burden 10^−2^ CFU/mL] or TTE has the outer bounds most influenced by *γ*_*s*_, consistent with clinical observations. In experimental regimens [**B** to **I**] PaMZ and BPaMZ regimens [**B** and **C**] sterilizing effect was faster than bactericidal effect, consistent with their mechanisms of action, so that TTE outer bounds were most influenced by *γ*_*f*_. In **D** to **E** the *γ*_*s*_ and *γ*_*f*_ values were similar, as shown by the parallel trajectories. In **F** to **I**, the *γ*_*s*_ and *γ*_*f*_ values were similar, but the slopes were steeper than standard therapy, so that TTE in the DBOS, DBO, DO, and BO were much shorter than for standard therapy. These regimens therefore have TTE that is not contingent just on sterilizing activity but also on activity against fast replicating bacilli.

### Translation of HFS-TB *γ*-slopes to patient sputum *γ*-slopes and therapy duration

What do these HFS-TB *γ*_*f*_ and *γ*_*s*_ slopes and TTE distributions translate to as regards to therapy duration in patients? To answer that, we converted the HFS-TB *γ*_*f*_ and *γ*_*s*_ slopes for each regimen to patients using our non-linear transformation factors and Latin hypercube sampling, with predicted patient *γ*_*f*_ and *γ*_*s*_ slope results shown in **Figure 7**. As a validation step for model predictions, we compared the model-predicted *γ*_*f*_ and *γ*_*s*_ for standard therapy in **Figure 7** to the *γ*_*f*_ of 0.86 [0.32-0.99] and *γ*_*s*_ of 0.19 [0.12-0.50] observed in patients’ sputa in two prior clinical trials.^20,21,34^ **Figure 7** shows virtually similar slopes for standard therapy based on our translation as observed in the clinic, which means that our quantitative translation was successful.

**Figure 7.**
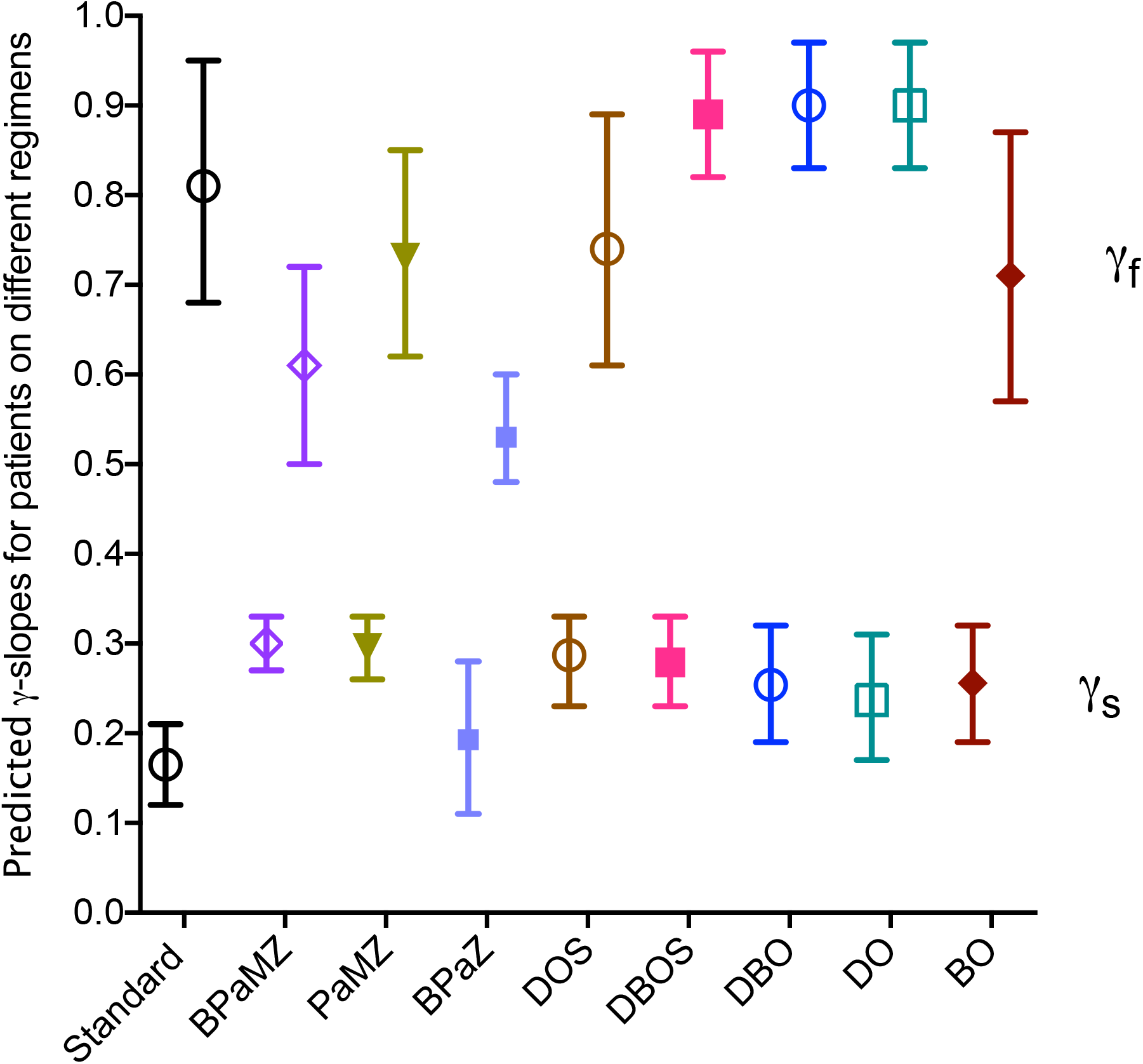
Predicted *γ*_*f*_ and *γ*_*s*_ slopes in sputa of 1,000 patients on different experimental regimens and standard therapy derived from HFS-TB. Symbols are mean slopes and error bars are 95% credible intervals. Data for the antagonistic delamanid-bedaquiline regimen and non-treated controls [with negative slopes] were omitted from graph for clarity purposes. The predicted *γ*_*f*_ slopes, which are generally higher that *γ*_*f*_, are shown in the upper rows and the predicted *γ*_*s*_ slopes in the lower rows. DO, DBO, and DBOS had high predicted *γ*_*f*_ slopes while BPaMZ overlapped with standard therapy, and the same regimens also had predicted *γ*_*s*_ slopes whose credible intervals overlapped with BPaMZ, which had the best *γ*_*s*_ slopes.

Figure 8. shows the predicted TTE of the entire bacterial population in patients, which captures both *γ*_*s*_ and *γ*_*f*_ and any antagonism or additivity thereof, and variability in initial bacterial burden in patients. Since predicted TTE in patients is a composite result of all these factors, we propose it as a better pharmacodynamic outcome for ranking therapy duration. **Figures 7** and **8**, based on *γ*_*s*_*-slope* thresholds and initial bacterial burden rule we derived in the clinic,^34^ show at least 5 regimens with potential to shorten therapy duration.

Figure 9. shows the TTE distribution predicted in patients converted into proportion of patients who achieved *Mtb* population extinction at 4 weeks, 6 weeks, 12 weeks, 16 weeks, and 6 months. Since the TTE equals the minimum duration of therapy required for relapse-free cure, **Figure 9** represents curves of time versus proportion of patients achieving relapse-free cure. We then ranked regimens based on the predicted minimum duration of therapy required for relapse-free cure in >95% of patients, which supersedes the rankings in **Table 1**.

In order to validate that the quantitative forecasting had been accurate we examined two aspects. First, **Figure 9** shows that standard therapy is predicted to achieve relapse-free cure in 85% of patients at 4 months and in >95% at 6 months, consistent with observations in clinical practice. Second, since the first 8 weeks data for BPaMZ and BPaZ were recently published after our sputum outcome predictions were presented to the sponsor,^55^ we compared the sputum clinical outcomes to our forecasts. In our model predictions, compared to standard therapy, the log-rank hazard ratio for time-to-extinction in BPaMZ was 2.98 [95% CI: 2.15-4.12] while that for BPaZ was 0.78 [95% CI: 0.60-0.99]. The hazard ratio for time to negative sputum in the clinical trial was reported as 3.3 [95% CI: 2.1-5.2] in MGIT culture or 2.3 [95% CI:1.5-3.4] on agar culture for BPaMZ, and 2.0 [95% CI: 1.3-3.2] in liquid cultures or 1.1 [95% CI:0.8-1.6] on agar for BPaZ.^55^ This means our forecasting step from HFS-TB to patients was successful.

**Figure 8.**
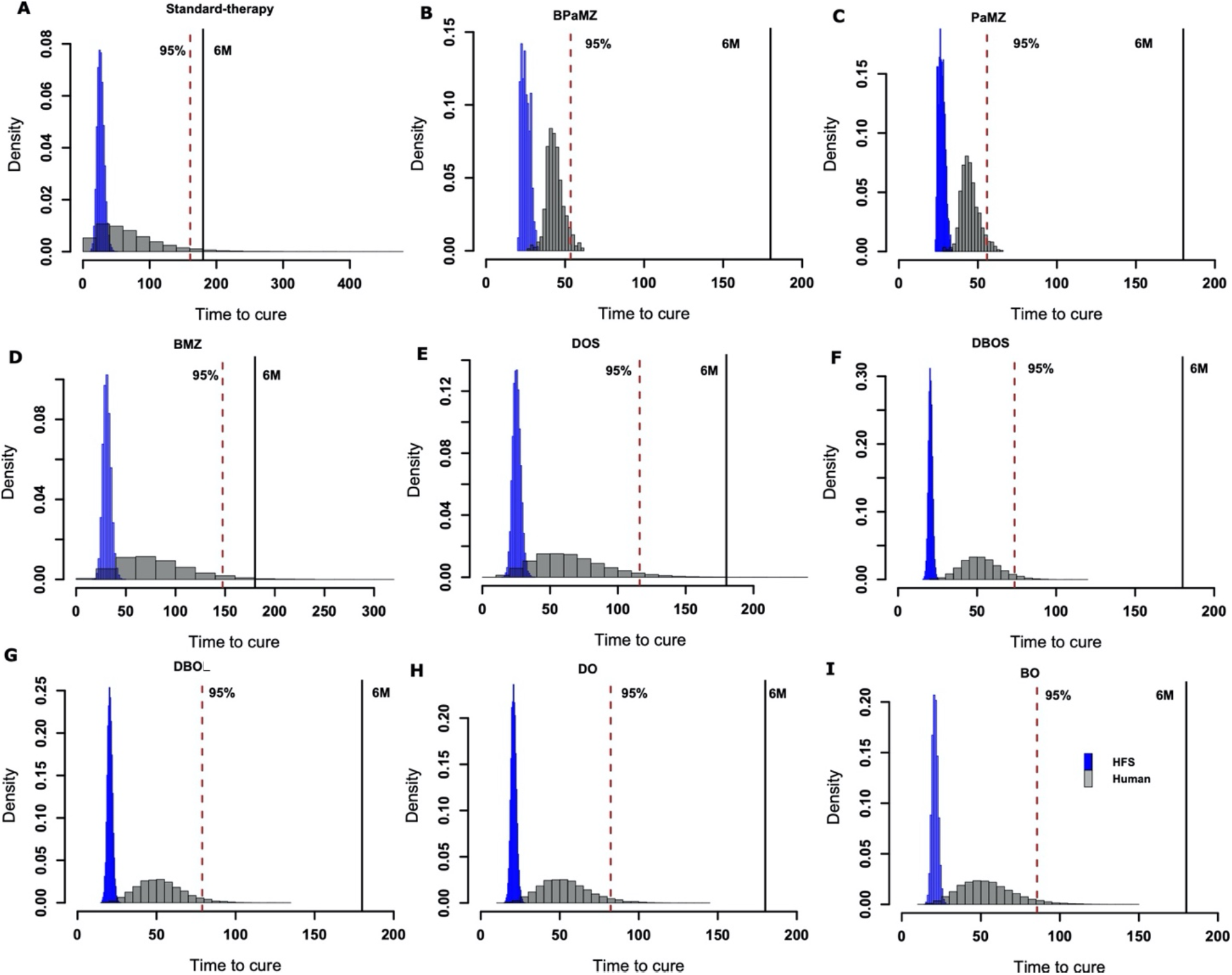
Time-to-extinction transformation from the hollow fiber to patients. HFS-TB time-to-extinction [TTE] results [blue] were translated to 1,000 patients’ groups [gray], leading to the distributions of TTE shown. It should be noted that the range of X-axis values [time in days] for standard therapy is 400 days while that BPaZ is 300 days, as opposed to the rest which are 0-200 days. The 6-months [180 days] TTE [black vertical line], the minimum duration required for cure [i.e., therapy duration], and 95% proportion of patients [red dotted vertical line] are shown for each regimen. (**A**) The standard regimen, 95% of patients would be cured at 160 days [23 weeks], which is what is observed in routine clinical care. (**B**-**I**) In the experimental regimens, (B) optimized BPaMZ and (F) DBOS, show 95% proportion with 8 weeks of therapy.

**Figure 9.**
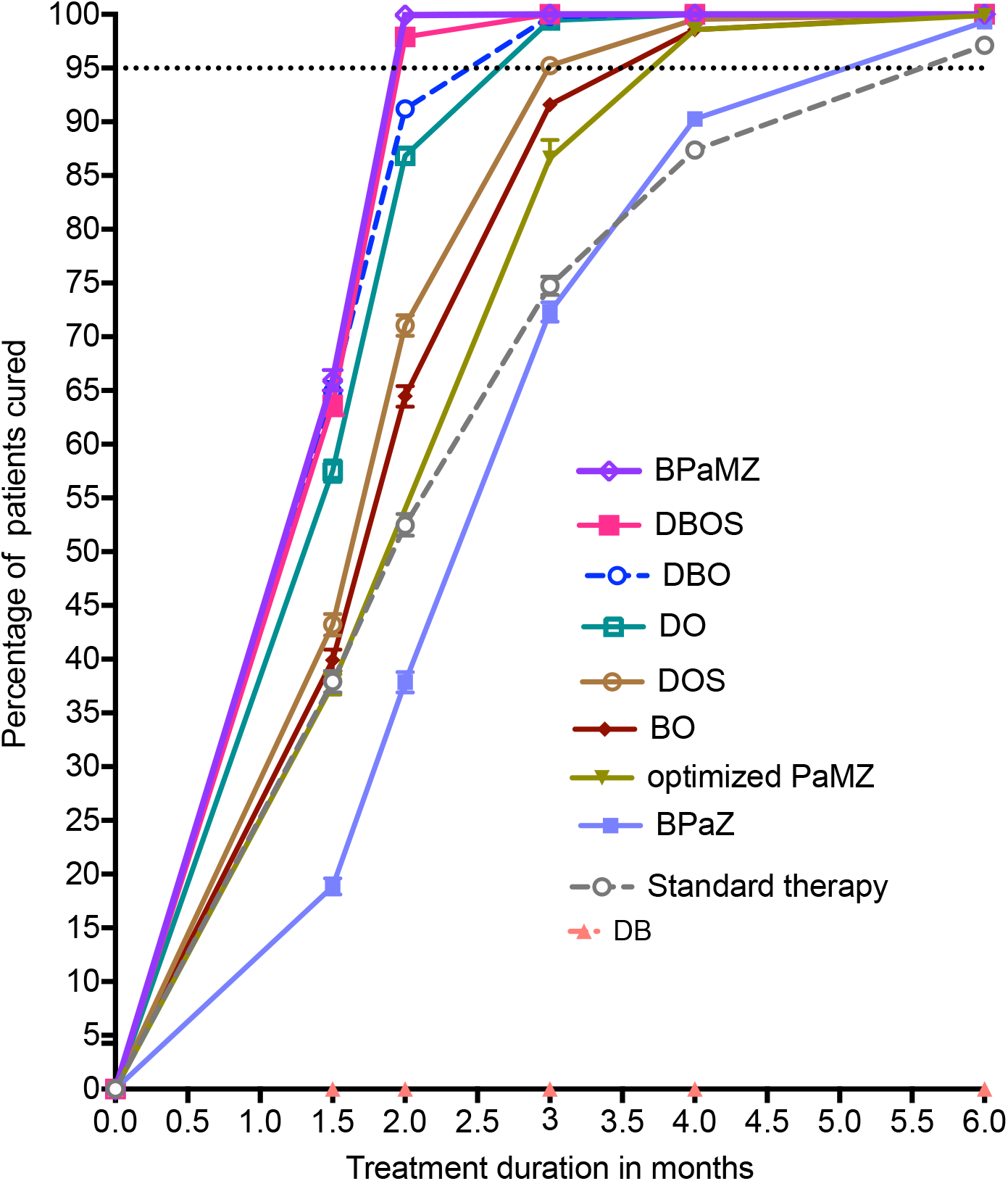
Proportion of 1,000 patients achieving extinction versus therapy duration. The duration of therapy that puts a minimum proportion of patients at risk of relapse or therapy failure is defined as the shortest time in which >95% of patients achieve *Mtb* population bacterial extinction, and hence relapse-free cure. Starting on the left, the BPaMZ [with PK/PD optimized PaMZ portion and standard dose bedaquiline] and DBOS ranked highest since they achieve this at 2 months, closely followed by DBO. BPaZ did not differ from standard therapy.

Figure 9. shows that DBOS and BPaMZ were predicted to achieve relapse-free cure with 8-weeks therapy duration in >95% of patients. Statistically, DBO was not much different if only slightly behind. In the second-best cluster [3-months duration], DO was significantly faster than DOS, but DBO was faster than DO, while DBOS [highest ranked cluster] was much faster than all three.

### A clinical trial strategy for testing clinical effectiveness within 2 years

Our predictions on DBOS, DBO and BPaMZ will need to be proven in clinical trials. Based on clinical data we have identified sputum *γ-*slopes during the first 8 weeks of therapy as the biomarkers of relapse-free outcomes up to 2 years later.^34^ We created a prediction rule of *γ-*slopes versus for minimum required duration of therapy; patients with slopes below thresholds in the biomarker had 8.25-fold higher risk of failure.^34^ The predicted patient sputum *Mtb γ-*slopes in **Figure 7** were taken as Bayesian priors as well as target slopes to be achieved in clinical trials,^34^ which allowed us a novel clinical trial design that puts a minimum number of patients at risk while expediting the process from decades to a few months. The DBOS mean composite *γ-*slopes, designated *m*_1_, and that for standard therapy [*m*_*2*_], were used to compute sample size based on Cohen’s *d* statistic to estimate the effect size with the Student’s two-sample t-test:

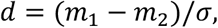

using the stringent thresholds for both sensitivity and specificity, for any diagnostic recommended by WHO, and the assumptions on unforeseen drug adverse events and attrition rates detailed in online methods. For a pooled standard deviation, *σ* =0.1, derived from both the standard arm and the experimental arms, an α set at 0.05, and (1-β) power of 0.8, assuming attrition [drop-out] rate of 10%, we calculated a sample size of 20 patients per group. In this design 20 patients per arm would be randomized to standard therapy versus DBOS for 8 weeks in patients with drug susceptible TB, over an observation period of 8 weeks. Patients would have weekly sputa for two months, and if the experimental regimens indeed achieved the target *γ-*slopes required to cure patients in 2-months shown in **Figure 9**, then the regimens will have a greater chance of achieving relapse-free cure in the predicted duration of therapy. The adverse events profile of the combination could also be compared to that of standard therapy over the two months. The follow up study would be an open-labeled clinical trial conducted of 120 patients with drug-susceptible TB who are treated for 2-months, and then followed regularly for 18 months to document relapse. Only patients with DS-TB should participate because if patients fail the experimental regimen they can be started on standard therapy, minimizing risk to patients, and optimizing risk-benefit ratios.

## DISCUSSION

First, the results generated here support DBOS as an ultrashort first-line pan-TB regimens, at doses that can be tolerated by patients. The diverse antibacterial mechanisms could explain the synergy: oxazolidinones inhibit bacterial protein synthesis, bedaquiline inhibits ATP production, the decaprenylphosphoryl-β-d-ribose 2’-epimerase (DprE1) inhibitor OPC-167832 inhibits cell wall synthesis, and delamanid inhibits the synthesis of the essential cell wall components methoxy-mycolic and keto-mycolic acid.^8,10,56^ However, the DBO regimen was not far behind the DBOS, with >90% cure predicted with 8 weeks therapy, and could also be ranked as first line, especially given the risk of oxazolidinone toxicity. In the 3-7% of patients predicted to fail DBOS or DBO, we propose derivation of a backup regimen with cure rates >95% in eight weeks so that in tandem we could cumulatively cure 99.91% of patients in four months. Thus ideally, the pan-TB regimen would not be a single regimen but consist of a first-line regimen plus a backup regimen composed of drugs that have alternative mechanisms of action. Importantly, if only novel drugs are used in the experimental regimens, and only patients susceptible to the current standard regimen are included in the studies, the current standard regimen will be a safe and reliable rescue treatment for those failing on the experimental arms.

Second, the identification of kill slopes predicted in patients on treatment shortening regimens, and the relationship between target kill slopes and duration of therapy in patients, allow for the design of novel TB clinical trials that require minimum number of patients to be put at risk of failure and death, and with a shorter timeline.^34^ Patients with DS-TB could be randomized to the DBOS or DBO regimen or to the standard first-line regimen over 8 weeks of therapy, and weekly liquid cultures with time-to-positivity over 8 weeks. At the end of 8 weeks, target criteria for the experimental arm are [1] slopes that indicate achievement of bacterial extinction at 8 weeks (i.e., test regimen meets target slopes), and [2] have at least two consecutive negative sputa, and [3] resolution of symptoms and [4] a lower or equivalent side effect profile to standard therapy. Since the outcome is driven by the bacteria kill slopes, themselves based on repetitive sampling [weekly] of each patient’s sputum, the sample size expected is only 20 patients per arm, greatly reducing patients at risk from several thousand with current practice. If the 8-week regimen and experiment meets target criteria, then a simple non-randomized cohort of about 120 patients in whom sputum is not tested for drug resistance would be treated with the 8-weeks regimen in a phase III open labelled trial, and then followed for 18 months for relapse.

Third, the finding that rapid microbial kill of fast-growing *Mtb* might be as important as slow-growing *Mtb*, especially as therapy duration gets shorter, has major implications on TB regimen design. The greater importance placed in eradication of slow/NRP bacteria as the “rate limiting step” was based on observations with current standard therapy, for which microbial kill of fast-growing *Mtb* is relatively rapid on a scale of few weeks but slow/NRP bacteria take half-a-year which made them the “rate limiting step”.^33,54^ This is based on the mechanisms of effect of rifampin, isoniazid and pyrazinamide, and the general refractoriness of slow-growing/NRP *Mtb* to these three drugs. However, such refractoriness, even for NRP, is not a universal rule, and is in part dependent on the mechanism of effect of specific first-line drugs. Here, we found that as durations of therapy approached 2 months, the time required to kill fast replicating *Mtb* also became a limiting factor.

Fourth, while we do not recommend two-drug combination regimens in the clinic, they were instrumental for design purposes. We found that some of the observed two-drug antagonism [DB] do not carry through when 3-or 4-drug combinations were used [DBO, DBOS]. The mechanistic basis of the two-drug pharmacodynamic antagonism are unclear, as is the reason why a third drug completely alters that. The possibility of effect of the culture media and Tween on bedaquiline in the BD dual therapy is unlikely since the BO performed dramatically well and better than standard therapy. Moreover, Nuermberger and colleagues have also demonstrated antagonism of the sister nitroimidazole [pretomanid] and bedaquiline in vivo in the mouse model of TB.^57^ These preclinical model data may also seem to be contradicted by clinical data that shows effectiveness of DB-based regimens. A systemic review of 87 patients treated with bedaquiline plus delamanid (47 concomitantly, 15 sequentially) found a sputum conversion rate of 81%; however the patients also received other concomitant anti-TB drugs and were not really on dual therapy.^58^ In a recent phase II clinical trial cumulative culture conversion by 24 weeks was 92% for bedaquiline, 91% for delamanid, and 95% for bedaquiline plus delamanid; however the patients were also multidrug background treatment, and not just the dual regimen.^59^ The DBO and DBOS regimens we recommend still have the BD-backbone, but the concomitant drugs made these effective regimens.

Finally, here we offer a platform that utilizes the HFS-TB to mimic each drug’s concentration-time profile that is encountered in patient lesions, followed by a quantitative translation pathway, to reduce the number of possible drug combinations to tractable numbers. The combination therapy HFS-TB drug doses and components were designed in May of 2019 by taking advantage of many of our HFS-TB monotherapy and dual therapy studies done in the prior few years, the HFS-TB studies reported here were then performed between June and August 2019 to final CFU/mL counts and TTE, and the modeling of *γ-*slopes and TTE were completed by end of August 2019, for a total of 4 months, and sent to sponsor. The repetitive sampling property of the HFS-TB is crucial for the identification of slopes and hence translation to patients for therapy duration. The platform output includes [1] the best drugs to be combined, [2] the best clinical doses to be combined, [3] the best dosing schedule to administer to each patient, [4] and crucially, the minimum duration of therapy, as well as [5] target slopes for use in design of expeditious trials. Thus, the platform not only helps design shorter therapy durations, but also shorter clinical development programs and pathways to adoptions of new regimens.

We acknowledge limitations of pre-clinical in vitro models, mathematical modeling approaches, and that we make several assumptions in our translational platform. Model accuracy using these assumptions has been good in the past, but this may not always continue to hold in in the future. Clinical trials will be the final arbiter. Second, the doses of pyrazinamide and moxifloxacin required for the BPaMZ regimen might be in the toxic range for humans. While standard doses of these drugs could work in a proportion of patients, these high doses would be required in the combination regimens for >95% of patients to be cured in 2 months. This could place many patients at high risk of adverse events. Third, some of the drugs we studied such as OPC-167832 have been administered for only limited durations of therapy, thus long-term adverse events when large populations of patients are treated at the specified doses are still unknown. Related to this is that several combinations have never been tested in humans, and thus we don’t know if they are safe. Fourth, and closely related, some combinations which could shorten therapy duration could be clinically imprudent from a toxicity standpoint. Thus, the early phase clinical trials we propose should also be designed in such a way as to maximize collection of early biomarkers of adverse events in patients on the combinations, so that therapy can be stopped before adverse events occur.

In summary, we demonstrate a useful *de novo* design approach of specific promising combination regimens from several thousand possibilities to a tractable number. In this HFS-TB-based head-to-head comparison of the nine regimens, repetitive sampling allowed for modelling bacterial kill trajectories of *Mtb* under treatment in different treatment refractoriness states, until bacterial population extinction. Category theory concepts were used to translate this HFS-TB dataset to that of patients’ slopes and TTE; repetitive sampling of patient sputum is the standard clinical approach. Duration of therapy was introduced as a new pharmacodynamic parameter to rank order regimens. Two regimens were predicted to cure patients in 8-weeks, and three DO-based three and four drug regimens predicted to cure with 12 weeks therapy duration. The role of slopes and TTE allowed us a new clinical trial design that uses HFS-TB-based slopes as Bayesian priors and requires only 20 patients per arm and de-risks clinical trials in terms of patients’ morbidity and mortality.

## METHODS

### Bacterial strain and supplies

*Mtb* strains and their culture conditions, as well as reagents used, and details of hollow fiber cartridges are discussed in detail in online methods.

### Minimum inhibitory concentrations

MICs were identified using 3 methods, agar dilution method, macrobroth dilution, and the MGIT, based on the Clinical and Laboratory Standards Institute guidelines and manufacturer instructions. Each MIC was performed twice.

### Hollow fiber study design

Novel regimens studies were carried out in the HFS-TB as described previously in the literature and in documents submitted to the EMA and US FDA.^2-5,11,12,18,19^ Full details of the HFS-TB design, concentration-time profiles used, bacterial growth parameters, and measurement of bacterial burden as well as drug concentrations, are in online supplementary methods

### Translation of Mtb bacterial burden trajectories in HFS-TB to human lungs

In human TB, *Mtb* is believed to exist as different phenotypic/metabolic states, in spatially disparate ecological heterogeneous micro-environments in different TB histopathological lesions.^20,33,34,54^ Further, different TB drugs exhibit different diffusion profiles based on dynamical sink gradients into these micro-environments to achieve different spatial drug exposures.^48^ Our modeling captures ecological interactions between different bacteria phenotypic/metabolic subpopulations (ii) the biology of their co-existence and selection for persistence (iii) how the introduction of antibiotics can induce extinction of all bacteria subpopulations in human lungs.^20^ The model is a system of coupled ODEs that track the fast replicating bacteria subpopulation and the hardest to kill subpopulation (to explain persistence and drug sterilization) at the site of infection.^20,34^ The

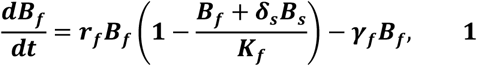

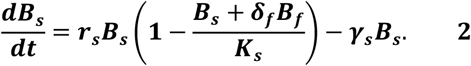

same equations were applied to sputa of patients undergoing repetitive sampling in clinical trials and routine TB care as well as to repetitive sampling in the HFS-TB, and the rate at which candidate TB regimens kill different bacterial subpopulations was evaluated, thereby estimating different bacteria subpopulations growth rates *r*_*s*_ and *r*_*f*_, and the corresponding drugs kill rates (*γ*_*s*_ and *γ*_*f*_), respectively.^20,34^ From these we calculated the time it took to extinguish all bacterial populations or time-to-extinction [TTE] in both the HFS-TB and patients’ lesions. The two datasets of kill rates and TTE form a structure-preserving map, as defined by category theory.^23-25^ Therefore, the elements of one can be mapped from another using a specified function. We have identified the composite and multi-step transformation function, ℱ: *H*_*F*_(= *F*(*H*^′^)), a *morphism*, which mapped the HFS-TB data to humans.^20^ Identifying this function and applying inverse modeling enabled data translation.

*Mtb* kill and growth suppression estimates, ***H***_***e***_ (= *H*_*e*_(*ε*_*f*_, *γ*_*f*_, *ε*_*s*_, *γ*_*s*_):, from the HFS-TB go through a transformation, ***T***_***f***_ (= *T*_*f*_ (*t*_ε *f*_, *t*_*γ f*_, *t*_ε *s*_, *t*_*γ s*_):, that map the HFS-TB estimates to the patient sputa estimates, (***C***_***e***_). Therefore, the transformation, ***C***_***e***_= ***T***_***f***_. ***H***_***e***_, gives the estimates that predict clinical results. The transformation factors, ***T***_***f***_, were estimated through fitting ***T***_***f***_. ***H***_***e***_ to the vitamin A clinical trial data in the past, and ***H***_***e***_ estimates were from the HFS-TB on standard therapy. The TTE was then calculated by simulating 1,000 samples using Latin Hypercube sampling.

### Monte Carlo experiments to translate from HFS-TB to clinical dosing

We used Monte Carlo experiments to identify clinical doses of the nine drugs studied that could achieve, or exceed, the AUC_0-24_/MIC attained in the HFS-TB, in 10,000 simulated subjects, using step-by-step procedures detailed in online supplementary methods.^7,36-47^

### A clinical trial strategy for testing clinical effectiveness within 2 years

Next, we computed sample size needed in phase II/III trials comparing new regimens to the standard 6-month regimen using the stringent thresholds for both sensitivity and specificity, for any diagnostic recommended by WHO. We made these four assumptions: 1) patients are randomized to regimens, and therefore, 2) initial bacterial burden would be similar between the regimens, 3) attrition rate of 20% comprising of a generous defaulting rate of 10% plus the unforeseen drug adverse events rates will be equal or less than the 10% encountered in recent BPaZ trial and 4) unequal variance in estimates of the *γ-*slopes. Therefore, we used pooled standard deviation, *σ*=0.1, which was derived from both the standard arm and the experimental arm, while α was set at 0.05 and (1-β) power was set at 0.8.

## Supporting information

Supplemental results and methods

## ONLINE MATERIAL

Supplementary results text

Table S1 and S2.

Figure S1, S2, and S3.

Supplementary methods

## ACKNOWLEDGEMENTS

Otsuka Pharmaceutical provided delamanid and OPC-167832.

## AUTHOR CONTRIBUTIONS

TG, DeH, DH: design of studies and experiments; MC, SS, DD: performance of HFS-TB experiments; GM: Mathematical modeling, translation of TB preclinical results to patients, therapy duration prediction; JGP: sample size calculation for clinical trials; RG: calculations of possible combinations based on Pascal and Fermat and PK/PD modeling for dose selection; AD, KD, TG, DH: new clinical trial designs using slopes. TG wrote first draft, and all authors had major editorial input to produce final draft.

