## Supplemental results and methods for "Novel tuberculosis combination regimens of two and three-months therapy duration"

**ONLINE MATERIAL**:

Supplementary results text

Table S1 and S2.

Figure S1, S2, and S3.

Supplementary methods

### Supplementary results text

#### Detailed Monte Carlo experiments output

We compared the Monte Carlo experiment [MCE] derived serum-based pharmacokinetic [PK] parameters at steady state to those achieved with the same dose in clinical studies performed by others in the clinic.[7,36-47] The MCE identified PK parameter estimates ±standard deviation at steady state such as total clearance [$CL_t$], central compartment volume [$V_C$], intercompartmental clearance [$CL_D$], peripheral volume [$V_p$], peak concentration [$C_{max}$] and $AUC_{0-24}$ achieved in the serum which we compared to those identified in actual clinical PK studies for validation purposes. The $AUC_{0-24}$s were then transformed into lung intralesional concentrations based on penetration ratios shown in the methods section.

In **Figure 3A,** an isoniazid dose of 600 mg/day MCE achieved a $CL_t$=20.72±3.3L, $V_C$=15.83±3.44L, $CL_D$=17.50±3.88L, $V_p$=22.11±2.66L;[37] the concentrations shown are double those encountered among slow acetylators in Tanzanian patients treated with half the dose [300 mg] reported to have a $C_{max}$=3.53 [IQR: 3.09; 3.83] mg/L and $AUC_{0-24}$=17.1 [IQR: 14.7; 21.2] mg*h/L.[45]

In **Figure 3B,** rifampin 35 mg/kg/day MCE achieved one compartment PK parameter estimates of $CL_t$=8.65±3.3L, and $V_C$=17.49±4.46L, and concentrations shown,[37] and the rifampin $C_{max}$= 21.6 [range:16.0–31.9] mg/L and $AUC_{0-24}$=113 [77.5–162] mg*h/L are similar to those encountered in patients taking rifampin 35 mg/kg in the study of Boeree et al.[39]

In **Figure 3C,** pyrazinamide 2,000 mg/day MCE achieved $CL_t$=3.84±0.88L, $V_C$=50.93±3.47L, and concentrations shown, similar to the $C_{max}$= 37.8 [IQR: 32.8; 44.5] mg/L and $AUC_{0-24}$=413 [IQR: 337; 546] mg*h/L encountered in patients in Tanzania and South Africa.[37,45]

In **Figure 3D** bedaquiline dosing simulated was 400 mg a day for 2 weeks, followed by 200 mg three times a week, thus the "steady" state period stretches out to 3 weeks, and at that time concentrations were similar to patients in the clinic in South Africa who achieved a $C_{max}$ of 4.07 [range:1.89-8.03] mg/L and an $AUC_{0-24}$ of 20.7 [range: 7.64-49] mg*h/L.[42] Bedaquiline is a 3 compartment model drug, with complex and delayed absorption, and the PK parameter estimates in the simulated subjected were a $CL_t$=2.62±1.02L, $V_C$=198±9L, $CL_2$=3.66±1.28L, $V_2$=8549±58L, $CL_3$=7.39±1.75L, $V_2$=2690±33L, similar to those encountered in patients.[41,42]

**In Figure 3E,** the pretomanid $CL_t$=3.28±1.42L, $V_C$=90.42±7.42L in the 10,000 subject MCE and the concentrations shown were similar to those in clinical studies. The $C_{max}$ achieved in patients in the clinic treated with 200 mg/day of 3.2 mg/L[46] was similar that achieved in in the MCE in **Figure 3E**.

In **Figure 3F** moxifloxacin, given at the high doses of 800 mg/day, and achieved $CL_t$=8.50±1.81L, $V_C$=114.1±4.62L, $CL_D$=2.89±0.74L, $V_p$=41.62±2.47L;[43] when co-administered with a rifamycin the 800 mg/day dose achieved a $C_{max}$=9.6 [range: 7.2-15.1] mg/L and $AUC_{0-24}$ of 78 [range: 53-120] mg*h/L in patients.[60]

In **Figure 3G,** the delamanid population pharmacokinetic parameter estimates were a LAG of 1.38±0.37 hrs, $CL_t$=37.33±8.78L, $V_C$=658.7±262.4L, $CL_D$=104.3±64.57L, and $V_p$=871.62±

341.9L.[43] In patients treated with delamanid 100 mg twice a day a $C_{max}$ of 0.4 [range: 0.3-0.5] mg/L and an $AUC_{0-24}$ of 9 [range: 6.3-10.7] mg/L are reported.[36]

In **Figure 3H,** the sutezolid PK parameter estimates in the MCEs were a $CL_t$=164.9±6.77L, $V_C$=12.1±10.7L, $CL_D$=32.43±2.2L,and $V_p$=258.9.34L; the concentrations in the simulated shown are virtually the same as the $C_{max}$=1.97±0.99 mg/L and $AUC_{0-24}$= 7.13±2.57 mg*h/L observed in patients on 1200 mg a day by Wallis et al.[7,47]

In **Figure 3I,** results are shwon for OPC-167832; the MCE pharmacokinetic parameter estimates of a $CL_t$=13.66±2.074L and V=303.1±9.771L were in the range of those seen in Multiple Ascending Dose/early bactericidal activity studies in patients with TB by Otsuka (unpublished; personal communications).

Overall, since the concentrations achieved in the blood in our MCE subjects were similar to those measured in the clinic, this means our simulation exercise was successful.

**Table S1. Regimens resulting from optimal design based on PK/PD principles**

| Regimen components | Name/designation |
|---|---|
| Sham treatment | Negative control |
| Isoniazid, rifampin, pyrazinamide | Standard therapy |
| Delamanid, bedaquiline* | DB |
| Delamanid, OPC-167832* | DO |
| Delamanid, bedaquiline, OPC-167832 | DBO |
| Delamanid, bedaquiline, OPC-167832, sutezolid | DBOS |
| Delamanid, OPC-167832, sutezolid | DOS |
| Optimized Pretomanid, moxifloxacin, pyrazinamide | Optimized PaMZ |
| Bedaquiline, pretomanid, moxifloxacin, pyrazinamide | BPaMZ |
| Bedaquiline, OPC-167832* | BO |
| Bedaquiline, pretomanid, pyrazinamide | BPaZ |

*The two-drug combination regimens are not intended for clinical use given concerns of drug-resistance, but for optimal experimental design purposes.

**Table S2. Slopes and time-to-extinction based on combined bactericidal and sterilizing effect hollow fiber system data**

| Parameter | Standard | BPaMZ | PaMZ | BPaZ | DOS | DBOS | DBO | DO | BO |
|---|---|---|---|---|---|---|---|---|---|
| $\gamma_f$ | 0.91(0.71-0.99) | 0.65(0.54-0.79) | 0.77(0.67-0.90) | 0.53(0.50-0.63) | 0.77(0.64-0.93) | 0.98(0.87-0.99) | 0.98(0.91-0.99) | 0.98(0.89-0.99) | 0.75(0.61-0.92) |
| $\gamma$s | 0.50(0.37-0.65) | 0.98(0.84-0.99) | 0.97(0.86-0.99) | 0.48(0.35-0.90) | 0.94(0.77-0.99) | 0.93(0.77-0.99) | 0.88(0.65-0.99) | .076(0.58-0.98) | 0.88(0.64-0.99) |
| k | 0.03(0.01-0.09) | 0.01(0.01-0.02 | 0.02(0.01-0.08) | 0.02(0.01-0.09) | 0.04(0.01-0.1) | 0.02(0.01-0.09) | 0.01(0.01-0.03) | 0.01(0.01-0.03) | 0.04(0.01-0.1) |
| TTE-HFS | 25.3(19.4-43.7 | 22.2(18.4-29.1) | 19.2(16.5-24.4) | 28.2(24.2-45.2) | 19.8 (16.4-25.6) | 15.3(15.2-18.8) | 15 (14.9-20) | 15.6(15.1-22.9) | 20.4(16.9-27) |

DB not shown because did not achieve extinction

**Figure S1. Growth of slow and fast replicating bacilli in non-treated controls and delamanid-bedaquiline dual combination.**

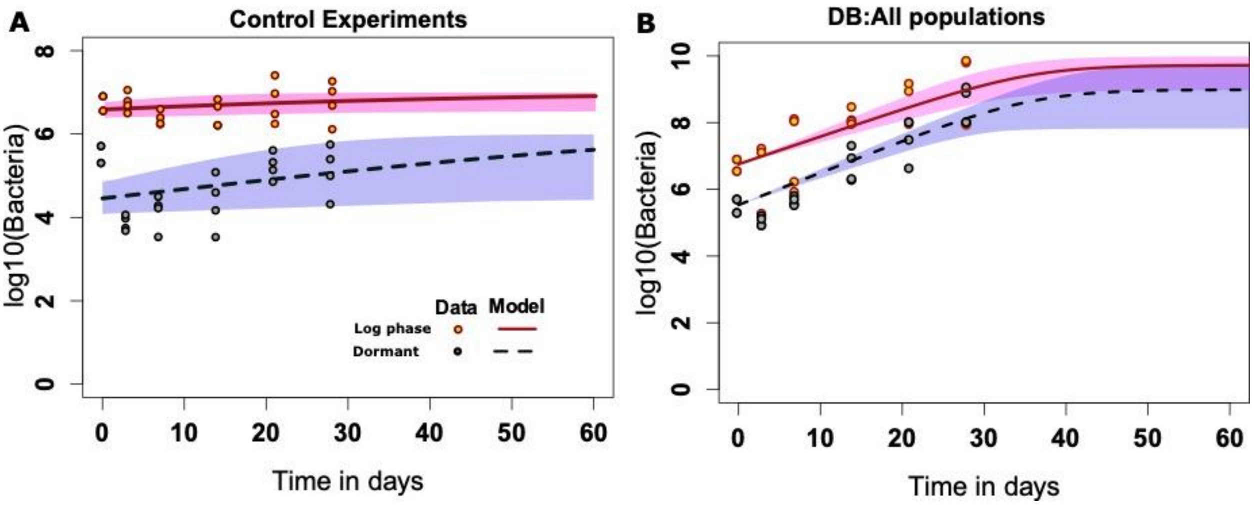


Grey dots represent dormant bacteria while orange dots represent fast replicating bacteria. The red line is the best model fit line for the fast replicating bacteria while the dotted black line is for the dormant subpopulation. The shaded regions represent the 95%CIs uncertainty in model to data fitting. Panel A shows the *Mycobacterium tuberculosis* control experiment data while Panel B shows the bedaquiline and delamanid combination therapy which is shown to fail.

### SUPPLEMENTARY METHODS

#### Bacterial strain and supplies.

*Mtb* H37Ra (ATCC #25177) and *Mtb* H37Rv (ATCC#27794) cultures were grown in Middlebrook 7H9 broth supplemented with 10% oleic acid, albumin, dextrose, and catalase (OADC) under shaking conditions at pH 6.8 or 5.8 for log phase and semi-dormant growth phases respectively. Sutezolid (PNU 100840) was purchased either from BOC Biosciences (Cas# 168828-58-8) or Sigma (#PZ0035), bedaquiline and pretomanid were purchased from BOC Biosciences; delamanid and OPC-167832 were received as gift from Otsuka Pharmaceutical; Pyrazinamide, isoniazid and rifampin were purchased from Sigma and Moxifloxacin was purchased from Baylor Pharmacy. Hollow fiber cartridges were purchased from FiberCell (Frederick, MD), BACTEC960 instrument and the Mycobacteria Growth Indicator Tubes (MGIT) were purchased from Becton Dickinson (Franklin Lakes, NJ).

#### Hollow fiber study design.

The HFS-TB units have been adapted for OPC-167832 and delamanid human-like concentration-time profiles, as described in detail in recent publications.[2] For bedaquiline containing regimen, we custom-designed a hybrid of polystyrene -based parts of the hollow fiber system, including central reservoirs/bottles and tubing while the hollow fibers were made of polyvinylidene difluoride. In addition, the non-specific binding of this drug was further overcome by administering bedaquiline at higher concentrations to account for further binding, based on earlier pharmacokinetic optimization studies and HFS-TB bedaquiline monotherapy studies. Thus, in the combination therapy systems that involved bedaquiline we used the hybrid system. Each of these systems had standard therapy and non-treated controls experimental conditions in preliminary studies, which were compared to the standard hollow fiber assembly, and they performed as expected and equivalent to standard hollow fiber assembly.

Target concentrations and pharmacokinetic parameters in the HFS-TB for each drug tested were as shown in the methods Table 1.

**Methods Table 1. MICs and target concentrations used in HFS-TB studies**

| Regimen | MIC mg/L | *Peak (mg/L) | $AUC_{0-24}$ (mg.h/L) | Half-life | Dose Justification |
|---|---|---|---|---|---|
| Isoniazid | 0.0625 | 6.8 | 24 | 3 | Optimized dose |
| Rifampin | 0.0625 | 6 | 22 | 3 | Optimized dose |
| Pyrazinamide | 25 | 54 | 1170 | 9 | Optimized dose |
| Moxifloxacin | 0.125 | 8.4 | 100 | 12 | $EC_{80}$ and resistance suppression [prior HFS] |
| Pretomanid | 0.01 | 0.25 | 5.73 | 18 | $EC_{80}$ [prior HFS] |
| Bedaquiline | 0.064 | 0.03 | 0.72 | 30 | Standard dose |
| Delamanid | 0.01 | 0.04 | 0.8 | 30 | $EC_{50}$ [additivity in HFS] |
| OPC167832 | 0.0005 | 0.22 | 4.03 | 30 | $EC_{80}$ [additivity in HFS] |
| Sutezolid | 0.25 | 4.5 | 16 | 4 | $EC_{80}$ [prior HFS] |

For HFS-TB-fast and HFS-TB-slow studies, *Mtb* culture in logarithmic growth phase or under acidic conditions [pH 5.8] were adjusted to ~6.0 $\log_{10}$ CFU/mL and 20 mL inoculum was inoculated in each of the peripheral compartment of HFS-TB. Since most of the drugs were AUC driven, they were infused in the central compartment everyday over 1 hr (standard therapy) or 4 hr (novel regimens) to achieve the intended AUCs. The concentration-time profiles of the drugs achieved in each of the HFS-TB was determined by sampling the central compartments over a period of 72-hours after the drug infusion based on optimal sampling theory. For quantitation of the total *Mtb* population, we sampled the peripheral compartment of each HFS-TB on day 0, 3, 7, 14, 21 and 28. Samples were centrifuged, supernatant was removed, washed twice with 0.9% saline and re-suspended in saline.  Samples thus prepared were then serially 10-fold diluted and spread on Middlebrook 7H10 agar supplemented with 10% OADC. The inoculated plates were incubated at 37°C for 21 days before the colonies were counted and expressed as colony forming units (CFU) per mL. For determining time to positivity (TTP), MGITs were labelled, 800 µL of OADC and 500 µL of sample was added and incubated in the BACTEC960 and fluorescent development was measured every hour until flags positive or up to 43 days.

**Drug concentration measurement assays.**

The assays we used to measure concentrations of isoniazid, rifampin, pyrazinamide, and moxifloxacin have been published in the past.[61,62] For delamanid, OPC, and pretomanid, LC-MS/MS analysis was performed using Waters Acquity UPLC coupled with Waters Xevo TQ mass spectrometer. Separation was achieved by injecting 10 µL of sample on a Waters Acquity UPLC HSS T3 column (50 x 2.1 mm; 1.8 µm) using a binary gradient. Solvents for UPLC were: (A) 0.1% aqueous formic acid, and (B) 0.1% formic acid in methanol. Samples were diluted 1:10 with internal standard solution containing pretomanid. The transitions used were *m/z* 457 to 176 (OPC), *m/z* 360 to 175 (pretomanid), and m/z 535.2 to 352 for delamanid. The between day percentage coefficient of variation (%CV) for analysis of low and high (brackets) quality controls were 1% (1%). The intraday %CV were 3% (2%). The lower limit of quantitation was 0.0025 µg/ml.

**Monte Carlo experiments to translate from HFS-TB to clinical dosing**.

Monte Carlo experiments [MCE] have been used for dose finding in TB by integrating these factors and HFS-TB output since 2002; they have a predictive accuracy of >94%.[16] We used the MCE to identify clinical doses of the nine drugs studied that could achieve, or exceed, the $AUC_{0-24}$/MIC attained in the HFS-TB, in 10,000 simulated subjects. PK parameter estimates and between patient variability [as %CV] that were used as domain of input in subroutine PRIOR of ADAPT software [University of Southern California] were shown in Methods **Table 2** below; these were from some of our own prior clinical studies as well as from the literature,[2,7,10,36,37,39,41-50] while those for OPC-167832 were from Otsuka Pharmaceutical Co. The sutezolid population pharmacokinetic model has not been published in full, but was presented as an oral presentation at the 7th Workshop on Clinical Pharmacology of Tuberculosis Drugs.[63] Data intralesional penetration $AUC_{0-24}$ ratios of rifampin, isoniazid, moxifloxacin, and pyrazinamide, were from our studies, and were those from the most difficult to reach part of the cavity, the air/caseum interface which formed a dynamical sink.[48,49] Drug intralesional concentrations of OPC-167832, delamanid, and bedaquiline used were assumed to be similar to those from animal studies.[2,10,50] For sutezolid we assumed similar penetration ratio to linezolid.[64,65] The pretomanid penetration ratio is unpublished: therefore, we examined different penetration ratio of up to 1, 2, and 3-fold, in the same range as the other nitroimidazole, delamanid.

**Methods Table 2. Population pharmacokinetic mean parameter estimates, and between-patient variability used in MCE**

| | Ka $hr^{-1}$ [%CV] | Total clearance L/hr [%CV] | Central volume L [%CV] | $CL_D$ L/hr [%CV] | Peripheral volume L [%CV] | Tlag hr [%CV] | Lung lesion-to-serum AUC ratio |
|---|---|---|---|---|---|---|---|
| Bedaquiline | 0.8 [40] | 2.62 [40.7] | 198 [43.3] | 3.66 [44.5] | 8550 [40] | 0.92 [40] | 0.86 |
| Delamanid | 0.6 [70.1] | 37.1 [24.5] | 655 [38.7] | 104 [67.8] | 870 [38.7] | 1.38 [60] | 2.0 |
| Isoniazid | 0.97 [10.28] | 8.63 [38.24] | 15.8 [72.78] | 17.5 [86.28] | 22.1 [32.44] | - | 0.4 |
| Moxifloxacin | 1.85 | 8.50 [12.6] | 114 | 2.90 | 41.6 | - | 1.26 |
| OPC-167832 | 1.1 [40.0] | 13.65 [31.0] | 303 [33.0] | - | - | - | 2.0 |
| Pretomanid* | 1.38 [70.78] | 3.30 [61.79] | 90.4 [60.9] | - | - | - | 1.0 or 2.0 or 3.0 |
| Pyrazinamide | 7.04 [58.81] | 3.86 [21.09] | 50.9 [23.77] | - | - | - | 0.45 |
| Rifampin | 0.35 [14.45] | 20.8 [54.33] | 17.5 [115.43] | - | - | - | 0.30 |
| Sutezolid | 0.88 [6.38] | 165 [27.8] | 162 [71.1] | 32.4 [14.8] | 258 [33] | 0.85 [10.7] | 0.53 |

*Parameters used are in fed state and not fasted state and are those observed in the BPaMZ regimen.
Bedaquiline was a 3-compartment model that included $V_3$ of 2690 L, and a $Q_3$ of 7.34 $hr^{-1}$.

Moxifloxacin, delamanid, and bedaquiline MIC distributions used were from the WHO "Technical Report on critical concentrations for drug susceptibility testing of medicines used in the treatment of drug-resistant tuberculosis" [https://apps.who.int/iris/handle/10665/260470]. The rifampin, isoniazid, and pyrazinamide MIC distributions were from our own work.[66] The sutezolid MIC distributions were from the clinical PK/PD study of Wallis et al, which are a small number of isolates, with 6/7 isolates having MICs ≤0.0625 mg/L, however the MIC distribution was truncated at 0.0625 mg/L and number of isolates tested small.[7] The range of 0.0625-0.25 mg/L is similar the same one in 47 MDR/XDR/DS-TB isolates reported by Yip et al in Hong Kong.[67] The pretomanid MIC distributions have not been published; however, documents submitted to the FDA indicate an MIC range of 0.005-0.48 mg/L, examined here. OPC MIC distribution was recently published.[10]

Doses of different drugs were used to identify the PK parameters and variance in 10,000 patients. The population pharmacokinetics parameters and their variances achieved in the simulated subjects were compared to those in the domain of input. Where available, serum peak concentrations and $AUC_{0-24}$s from different PK studies than those used in the domain of input were compared to those in simulated subjects were used as an extra external validation step. The intralesional $AUC_{0-24}$ of each drug for each dose was used to identify the $AUC_{0-24}$/MIC ratio at each MIC in each of the 10,000 patients; the proportion of patients at each MIC was defined as the target attainment probability [TAP].
